# RNA-seq analysis of a zebrafish *tlr2* mutant shows a broad function of this Toll-like receptor in transcriptional and metabolic control and defense to *Mycobacterium marinum* infection

**DOI:** 10.1101/742601

**Authors:** Shuxin Yang, Wanbin Hu, Yasuhito Shimada, Magnus Münch, Rubén Marín-Juez, Annemarie H. Meijer, Herman P. Spaink

## Abstract

**Background:** The function of Toll-like receptor 2 (TLR2) in host defense against pathogens, especially *Mycobacterium tuberculosis* (Mtb) is poorly understood. To investigate the role of TLR2 during mycobacterial infection, we analyzed the response of *tlr2* zebrafish mutant larvae to infection with *Mycobacterium marinum* (Mm), a close relative to Mtb, as a model for tuberculosis. We measured infection burdens and transcriptome responses using RNA deep sequencing in mutant and control larvae.

**Results:** *tlr2* mutant embryos at 2 dpf do not show morphological alterations or differences in the number of macrophages and neutrophils when compared to control embryos. However, we found substantial changes in gene expression in these mutants, particularly in developmental and metabolic pathways, when compared with the heterozygote *tlr2*^+/−^ control. After Mm infection, bacterial burden was six to ten fold higher in *tlr2*^−/−^ larvae than in *tlr2*^+/−^, or *tlr2^+/+^* larvae, indicating that Tlr2 acts as a protective factor in zebrafish host defense. RNAseq analysis of infected *tlr2^−/−^* versus *tlr2^+/−^* shows that the number of up-regulated and down-regulated genes in response to infection was greatly diminished in *tlr2* mutants by at least 2 fold and 10 fold, respectively. Analysis of the transcriptome data and qPCR validation shows that Mm infection of *tlr2* mutants leads to decreased mRNA levels of genes involved in inflammation and immune responses, including *il1b*, *tnfb*, *cxcl11aa/ac*, *fosl1a*, and *cebpb*. Furthermore, RNAseq analyses revealed that the expression of genes for Maf family transcription factors, vitamin D receptors, and Dicps proteins is significantly altered in *tlr2* mutants with or without infection. In addition, the data indicate a function of Tlr2 in the control of induction of cytokines and chemokines, such as the CXCR3-CXCL11 signaling axis.

**Conclusion:** The transcriptome and infection burden analyses give support for a function of TLR2 in host defense against mycobacteria. Transcriptome analysis revealed *tlr2*-specific pathways involved in Mm infection, which are related to responses to Mtb infection in human macrophages. Considering its dominant function in control of transcriptional processes that govern defense responses and metabolism, the TLR2 protein can be expected to be also of importance for other infectious diseases and interactions with the microbiome.

## Background

*Mycobacterium tuberculosis* (Mtb) is the causative agent of tuberculosis (TB), which infects nearly 23% of the world’s population, and kills about 1.6 million people annually (WHO Global Tuberculosis Report 2018). TB is characterized by the formation of granulomas, aggregates of infected macrophages and other immune cells, not only in the lung but also in other tissues and organs[1]. The formation of granulomas is the result of a concerted action of host innate and adaptive immunity [2-4].

Innate immune responses play a critical role in defense against TB infection in the host, and for a major part these processes are mediated by Toll-like receptors (TLRs), a conserved family of pattern recognition receptors. TLR2 is one of the most widely reported members of the TLR family to be involved in defense against Mtb by virtue of its recognition of cell wall-associated components associated with this pathogen [5-7]. Following mycobacterial infection in human cell cultures, TLR2 dimerizes with TLR1 or TLR6, and recognizes mycobacterial components such as cell wall glycolipids LAM and LM [8], 38-kDa and 19-kDa mycobacterial glycoprotein (LpqH) [9-11], phosphatidylinositol mannoside (PIM) [12], and triacylated (TLR2/TLR1) [13, 14] or diacylated (TLR2/TLR6) lipoproteins [15, 16]. Then, these heterodimers recruit the MYD88 and TIRAP (MAL) proteins to activate the IRAK (1 and 4)/TRAF6/IKK (α or β) cascade, which subsequently leads to the ubiquitination of IκBα and the activation of transcription factor NF-κB or AP-1 to induce the expression of host defense genes and cytokine and chemokine responses [17, 18]. Once released after further processing, these cytokines and chemokines attract migration of macrophages and neutrophils to the infection site and activate the microbicidal functions of these cells.

TLR2 has been shown to be important for granuloma formation in Mtb infection in mouse infection model [19]. Other studies have reported that *Tlr2*^−/−^ mice lose control to high dose infection of Mtb delivered by aerosol administration and show higher susceptibility to Mtb infection compared to the wild type [20, 21]. However, using lower doses of infection of 100 bacteria, there are controversial results as to the function of TLR2 in defense to Mtb in rodent models [20-22]. Furthermore, the function of TLR2 in susceptibility to Mtb is still unclear because several independent studies have reported polymorphisms of TLR2 (TLR1 and TLR6) in humans that have been linked with TB susceptibility, whereas others have been unable to find such links [23, 24].

Several studies suggest that the interaction between TLR2 and Mtb or other pathogens does not always promote the killing of bacteria, but can in fact be part of the pathogens’s strategy to evade the immune system [25-27]. TLR2 has been reported to inhibit MHC-II expression on the surface of murine macrophages, thereby preventing presentation of Mtb antigens, which may allow intracellular Mtb to evade immune surveillance and maintain chronic infection [11, 28]. Furthermore, LprG from Mtb was reported to inhibit human macrophage class II MHC antigen processing through TLR signaling [25]. In other infection systems, *Tlr2* mutant mice have an increased resistance against infection of *Candida albicans* [29] and *Yersinia pestis* [30]. It has been proposed that this phenotype is the result of Tlr2-dependent induction of the anti-inflammatory cytokine IL-10 [30]. Furthermore, TLR2 activation inhibits the release of IL-12 via activation of the cFos transcription factor. It also has been shown that IFN-γ or IFN-γ-induced signals [31] are inhibited after infection of murine macrophages in a Tlr2-dependent fashion [32]. These effects show that TLR2 activation can yield a bias to T helper Type 2 (Th2) cells [33], and by breaking the Th1/Th2 balance can lead to less Th1 type responses and reducing the killing of intracellular pathogens. However, most of the molecular mechanisms underlying TLR2 functions remain unknown and a better understanding of the TLR2-mediated immune response and immune evasion can help in planning prevention and therapy strategies against Mtb infection.

Animal models have shown their power in studies of the mechanisms of interaction of host and TB pathogens, and discovering new anti-TB drugs. Zebrafish adult and larvae models have become a useful complement for rodent studies, for three important reasons. First, zebrafish have a 3-4 weeks separation stage between development of innate and adaptive immunity after fertilization [34, 35], which gives the possibility to study the host innate immune response to infection in the absence of adaptive immune responses. Second, zebrafish can be infected by *Mycobacterium marinum* (Mm), a natural pathogen of cold-blooded vertebrates and a close relative of Mtb, which can induce granuloma formation in zebrafish, similar to human TB [36, 37]. Third, the transparent larvae are ideal for imaging the early steps of the infection process in real time. Hence, zebrafish has earned its place of being a versatile tuberculosis model [1, 38, 39].

In our previous study, we demonstrated that the mammalian TLR2 ligand Pam3CSK4, a synthetic triacylated lipopeptide that mimics the triacylated lipoprotein of mycobacteria, could also specifically activate the zebrafish Tlr2 pathway, inducing *fosl1a* and *cebpb* gene upregulation [40]. In the current study, to further explore the involvement of Tlr2 in Mm infection, we conducted infection studies using *tlr2* mutant zebrafish. We found that a *tlr2* mutation led to an overall increased Mm bacterial burden, corresponding with reduced presence of macrophages in the granulomatous aggregates and more extracellular growth. These results indicate that Tlr2 plays an important role in protecting the host during the early stage of mycobacterial infection. In addition, we performed RNA deep sequencing (RNAseq) and determined a Tlr2-specific gene list for the response to Mm infection. This analysis revealed that most of transcriptional downregulation caused by Mm infection in control animals was abrogated by *tlr2* mutation, in addition to a dampening effect on the upregulation of transcription factors and inflammatory genes.

## Results

### 1. Characterisation of a *tlr2* mutant zebrafish line

The *tlr2*^sa19423^ mutant (*tlr2*^−/−^) carries a thymine to adenine point mutation that creates a premature stop codon (Fig. 1a), which is located in the sequence coding for the C-terminus of the leucine-rich repeat (LRR) domain. This leads to a truncated protein without the Toll/IL-1 receptor (TIR) domain, which is required for the interaction with Myd88 and Tirap (Mal) [41, 42].

**Figure 1.**
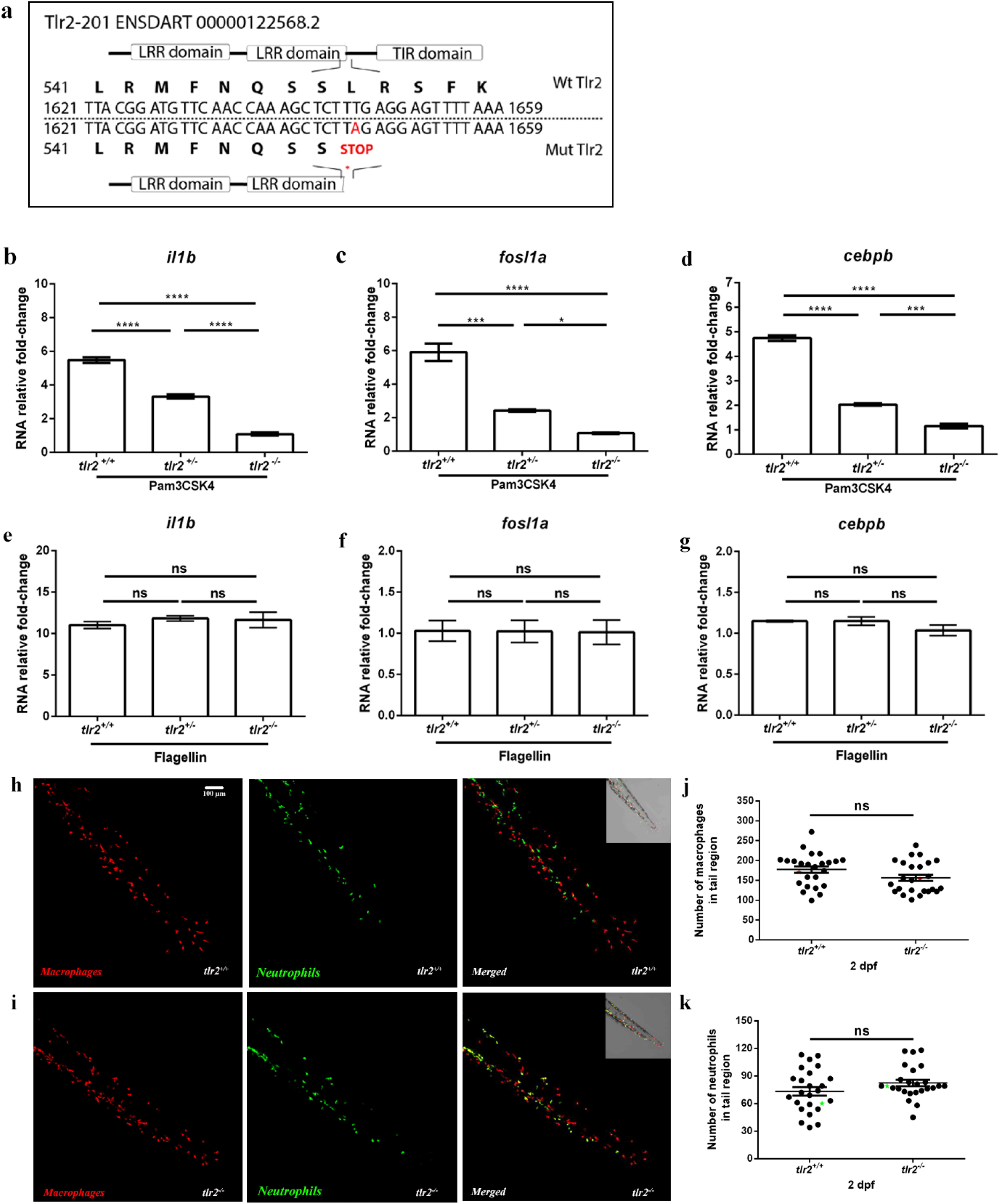
Characterization of the Tlr2 mutant. (a) mutant DNA and protein sequence. A point mutation (T to A) in the C-terminal of the second LRR domain of zebrafish Tlr2 introduces a premature stop codon. The predicted truncated protein lacks the whole TIR domain. Nucleotide and amino acid positions are indicated with respect to the translation start codon. *tlr2^+/+^*, *tlr2^+/−^* and *tlr2^−/−^* embryos were injected at 27 hpf with 1 ng Pam3CSK4 (b-d) or 0.1ng flagellin (e-g) and expression levels of *il1b*, *fosl1a* and *cebpb* were determined at 1hour post injection by qPCR. Data (mean ± SEM) are combined from three biological replicates (n=10 embryos per group) and expressed relative to their corresponding mock injection (water) control, which set at 1. Statistical significance of differences between mock and Pam3CSK4 groups was determined by one-way ANOVA with Tukey’s Multiple Comparison method as a post-hoc test (b-g), ns, non-significant; *, *P*<0.05; **, *P*<0.01; ***, *P*<0.001; ****, *P*<0.0001. (h, i). Representative Stereo images of the caudal hematopoietic tissue of *tlr2^+/+^ Tg*(*mpeg1:mCherry-F);TgBAC(mpx:EGFP)* (h) and *tlr2^−/−^ Tg*(*mpeg1:mCherry-F);TgBAC(mpx:EGFP)* (i) were taken at 2 dpf for quantification of macrophages and neutrophils numbers. At 2 dpf, numbers of mCherry-labeled macrophages (j) and GFP-labeled neutrophils (k) were counted using Leica TCS SP8 confocal laser scanning microscopy (CLSM) of transgenic lines. Data (mean ± SEM) were combined from two independent experiments. No significant differences (ns) in the number of macrophages (j) and neutrophils (k) was detected with a t-test.

To confirm whether *tlr2* mutation blocks its downstream pathway, we analyzed the gene expression profiles of zebrafish treated with the TLR2 agonist, Pam3CSK4 similarly as in our previous work [43], now also including the heterozygote mutant (*tlr2*^+/−^). Pam3CSK4 was injected into the blood island of zebrafish embryos at 27 hours post fertilization (hpf). One hour after injection (hpi), we collected samples and performed qPCR to analyze the expression levels of CCAAT/enhancer-binding protein beta (*cebpb*) and FOS Like Antigen 1a (*fosl1a*), previously shown to be specific targets of Tlr2 signaling [40]. In wild-type siblings and *tlr2^+/^* ^sa19423^ heterozygotes (*tlr2^+/^*^−^), the expression levels of *cebpb* and *fosl1a*, as well as the inflammatory gene *il1b* were significantly induced upon Pam3CSK4 injection, whereas *tlr2^−/−^* showed no significant response (Fig. 1b-d). *tlr2^+/^*^−^ animals showed a lower induction of these marker genes than the wild-types, indicating that there is an effect of the *tlr2* mutation even in the heterozygote. To confirm that these results are specific to the Tlr2 pathway, we injected flagellin, a Tlr5 agonist, into 27 hpf embryos. Flagellin induced *il1b* expression, but not *cebpb* and *fosl1a* expression in wild-type siblings, *tlr2^+/−^* and *tlr2^−/−^* larvae (Fig. 1e-g). Overall, these data show that *tlr2* mutation specifically blocks the Tlr2 downstream pathways.

To determine if immune cell development was affected by *tlr2* mutation, we used the double-transgenic line *tlr2^+/+^ Tg*(*mpeg1:mCherry-F);TgBAC(mpx:EGFP)* and *tlr2^−/−^* *Tg*(*mpeg1:mCherry-F);TgBAC(mpx:EGFP)* to count the number of macrophages and neutrophils at 2 dpf. The results show that there is no significant difference in the number of macrophages and neutrophils at 2 dpf between the wild type and mutant (Fig. 1j, k).

### 2 Comparison of gene expression profiles of *tlr2* homozygote and heterozygote mutants in the absence of infection

In order to investigate the systemic effects of the *tlr2* mutation we compared basal levels of gene expression in the absence of infection between *tlr2* homozygote and heterozygote mutants (Fig. 2). We chose to use the heterozygote *tlr2^+/−^* line as a control since these are genetically as comparable as possible, considering the fact that random ENU mutations could still contaminate our background even after 3 generations of outcrossing and the use of sibling lines. To further minimize the effect of polymorphisms, we pooled 10 larvae in each of our samples. These results show that there is a large group of genes that are expressed differently even at extremely stringent false discovery rate (FDR) adjusted *p-*value of 10^−10^ (Fig. 2). Since the results of the DEseq2 analyses showed such a surprisingly large number of significant differences we also used another statistical method for analyses, called edgeR, that differs in normalization and estimation of the dispersion parameters [44]. EdgeR analyses confirmed the statistical significance of the differences between the homozygote and heterozygote mutants (Fig. 2a and 2b). While no differences in numbers of mpeg-positive macrophages were detected (Fig. 1j, k), the RNAseq analysis showed that the *mpeg1* gene is expressed approximately 2 fold higher in the *tlr2*^+/−^ control than in the *tlr2*^−/−^ mutant (Supplementary Table 1). This was confirmed by pixel count measurements in the transgenic line *tlr2^+/+^ Tg*(*mpeg1:mCherry-F);TgBAC(mpx:EGFP)* and *tlr2^−/−^ Tg*(*mpeg1:mCherry-F);TgBAC(mpx:EGFP)* (Fig. S1 a, b). These results showed lower fluorescence of the mCherry *mpeg1* reporter as compared to the wild-type siblings at 2 dpf (Fig. S1 a) whereas no difference was detected for the eGFP *mpx* reporter (Fig. S1 b). Therefore these *in vivo* results confirm the RNAseq data. We also examined the expression levels of the *tlr2* gene in the homozygote and heterozygote mutants in the RNAseq data (Fig. S2). The results show that, although *tlr2* is very lowly expressed, there is no difference in expression levels between the homozygote and heterozygote mutants, indicating that there is no non-sense mediated mRNA decay (NMD).

**Figure 2.**
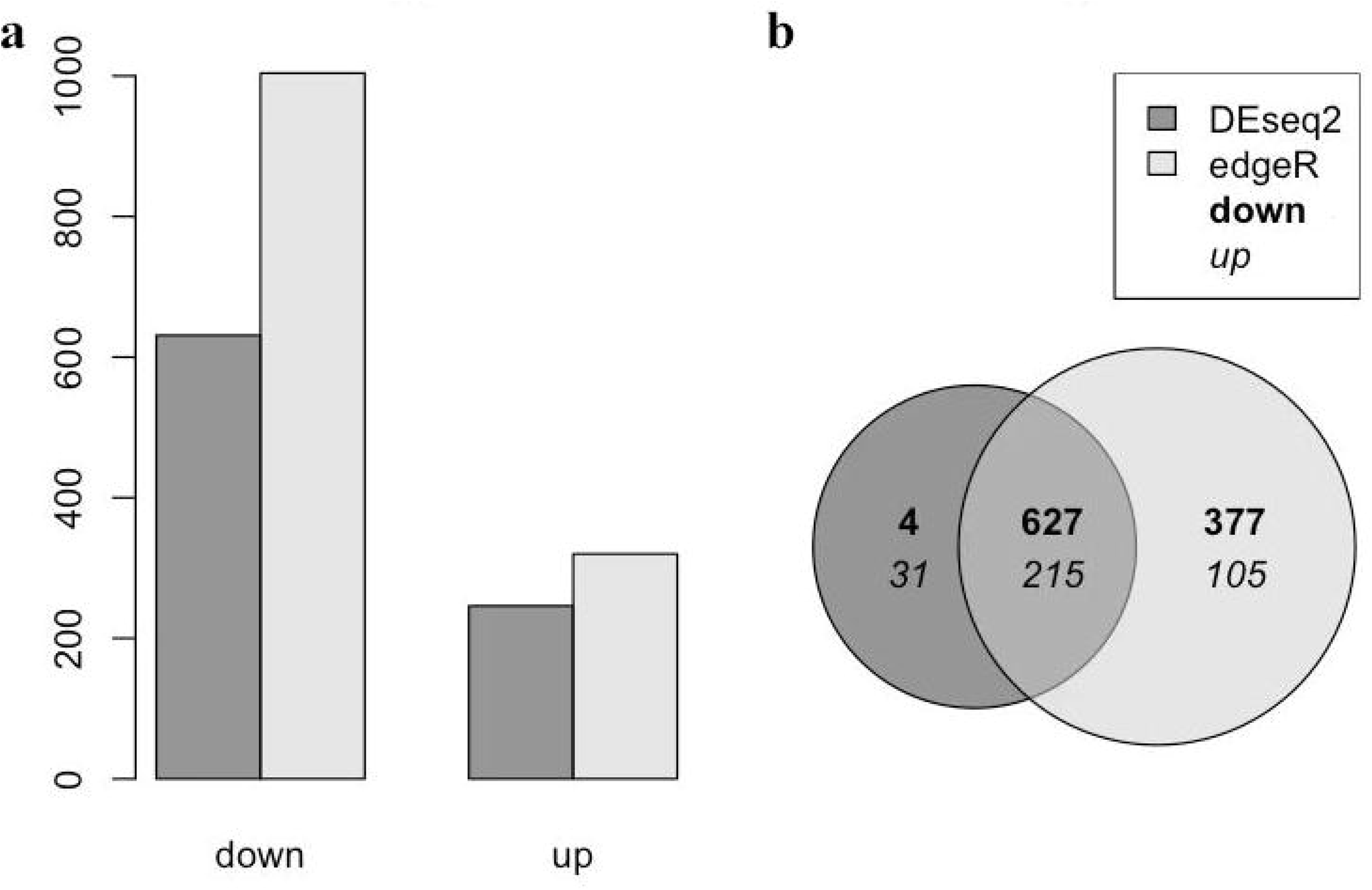
Gene expression in the absence of infection between Tlr2 mutants. (a) a number of down- and up-regulated genes per method (edgeR and DEseq2) and (b) a Venn diagram that compares the down-, up- and non-differentially expressed genes per method at a significance level of 10^−10^ in the edgeR and DEseq2 analyses methods. Down-regulated means that a gene is less expressed in the *tlr2*^+/−^ compared to the *tlr2*^−/−^ strain.

To investigate whether the basal expression level differences of genes might be relevant to the Tlr2 pathway, we performed GO analysis of a set of genes that were expressed differently with the very low FDR adjust *P* value of 10^−10^ (Supplementary Table 2). Gene ontology analyses of the gene sets that differ with an FDR of 10^−10^ in both analyses show that there is a large enrichment of genes belonging to the GO terms related to neural development (Supplementary Table 2). In addition, we have also analyzed differences with a fold change criterion of two and FDR of 0.05. These analyses indicated that under the GO category “Transcription factor genes” only two categories of genes including the c-Maf transcription factor were present (Supplementary Table 3). Pathway analysis shows that genes that function in glucose metabolism are differentially expressed in the *tlr2* homozygote versus heterozygote mutant (Fig. S3).

### 3 Tlr2-specific gene expression profiles after Mm infection

To study the role of Tlr2 in Mm infection we tested the *tlr2* mutant line in comparison with the heterozygote and wild-type sibling controls (Fig. 3). No differences in bacterial burden were observed at 3 dpi (Fig. 3d). However, we found that bacterial burden was significantly higher in *tlr2^−/−^* than in *tlr2^+/−^* and wild-type larvae at 4 dpi (Fig. 3e). There was no significant difference in bacterial infection burden between the heterozygote mutant strain (*tlr2^+/−^*) and the wild-type siblings (*tlr2^+/+^*).

**Figure 3.**
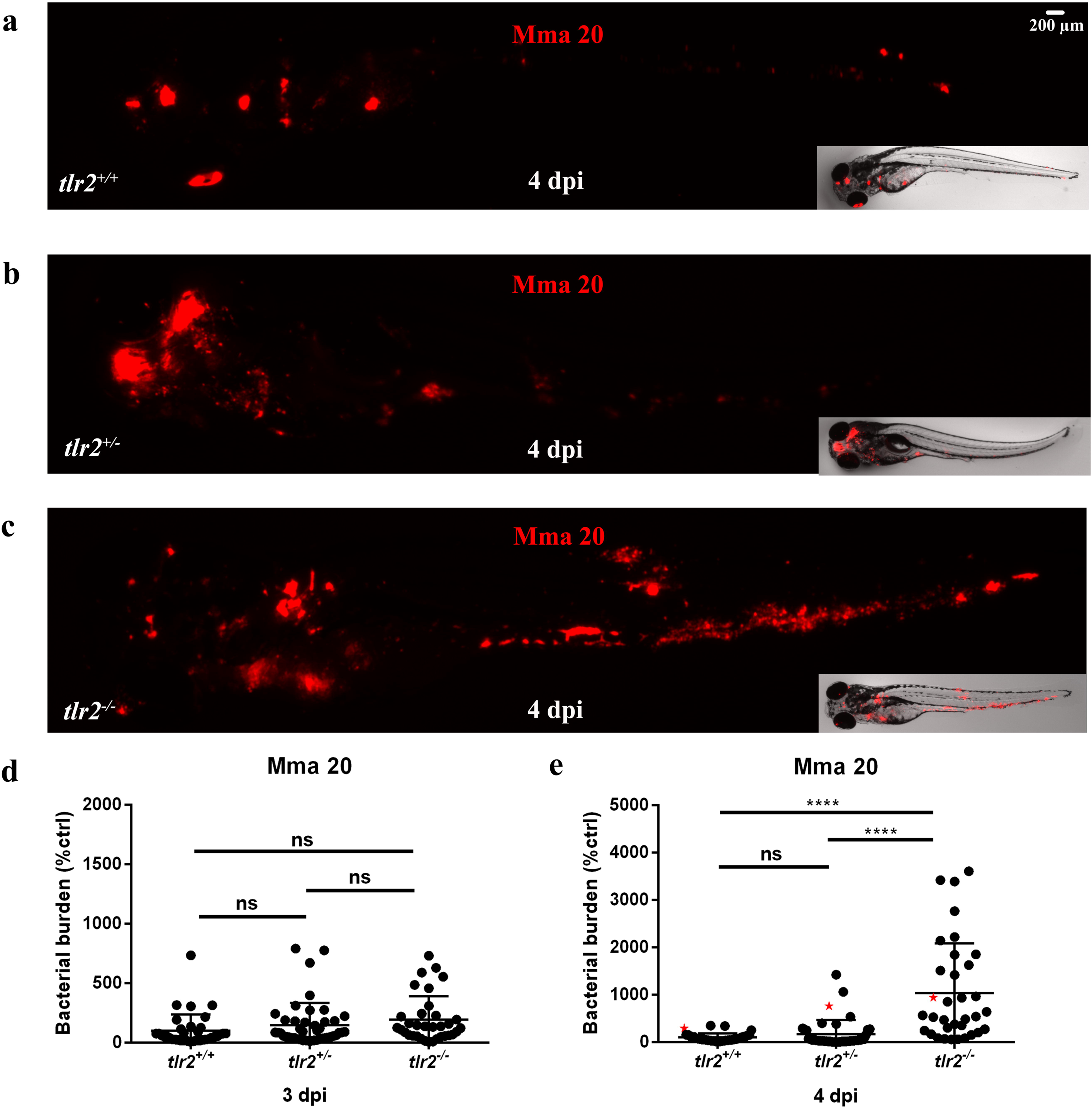
Quantification of bacterial burden. *tlr2^+/+^* (a), *tlr2^+/−^* (b) and *tlr2^−/−^* (c) embryos were infected with mCherry-labeled *M. marinum* strain Mma 20 at a dose of ∼150 CFU by caudal vein infection at 28 hpf. Representative images for bacterial pixel count in *tlr2^+/+^* (a), *tlr2^+/−^* (b) and *tlr2^−/−^* (c) were taken at 4 dpi. Bacterial burden of *tlr2^+/+^*, *tlr2^+/−^* and *tlr2^−/−^* were also quantified at 3 dpi (d) and 4 dpi (e). Bacterial burdens were quantified by using bacterial fluorescence pixels. Red stars in (e) indicate the data for the representative images shown in (a-c). In (d, e), data (mean ± SEM) were combined from two independent experiments. Statistical significance of differences was determined by one-way ANOVA with Tukey’s Multiple Comparison method as a post-hoc test for comparison between more than two groups (d, e). ns, non-significant; ***, *P*<0.001; ****, *P*<0.0001.

Next, we set out to assess the general inflammation and specific immune responses in *tlr2* mutants. We first studied by qPCR analysis the expression of *il1b*, *tnfa*, *tnfb*, *irg1l*, and *tlr2* specific transcription factors *fosl1a* and *cebpb* in *tlr2*^−/−^ and *tlr2*^+/−^ larvae upon Mm (strain Mma20) infection at 4 dpi. Our results show that the induction levels of *il1b*, *tnfb, fosl1a* and *cebpb* in *tlr2*^−/−^ larvae were significantly reduced when compared to the heterozygotes in the infected condition (Fig. 4). *tlr2*^−/−^ larvae failed to upregulate *fosl1a* and *cebpb* expression in response to Mm administration (Fig. 4e, f). In our previous work, we showed that Cxcl11-like chemokines play a crucial role in macrophage-mediated granuloma formation upon Mm infection [45, 46]. We therefore conducted qPCR to assess the expression levels of *cxcl11-like* genes, previously shown to be induced by infection [45], including *cxcl11aa* and *cxcl11ac* (Fig. 4g, h). The expression levels of *cxcl11aa* and *cxcl11ac* was significantly higher in *tlr2^+/−^* larvae than in *tlr2^−/−^* upon Mm infection (Fig. 4g, h). These results indicate that *tlr2* mutation results in a defective immune or inflammatory response to Mm infection.

**Figure 4.**
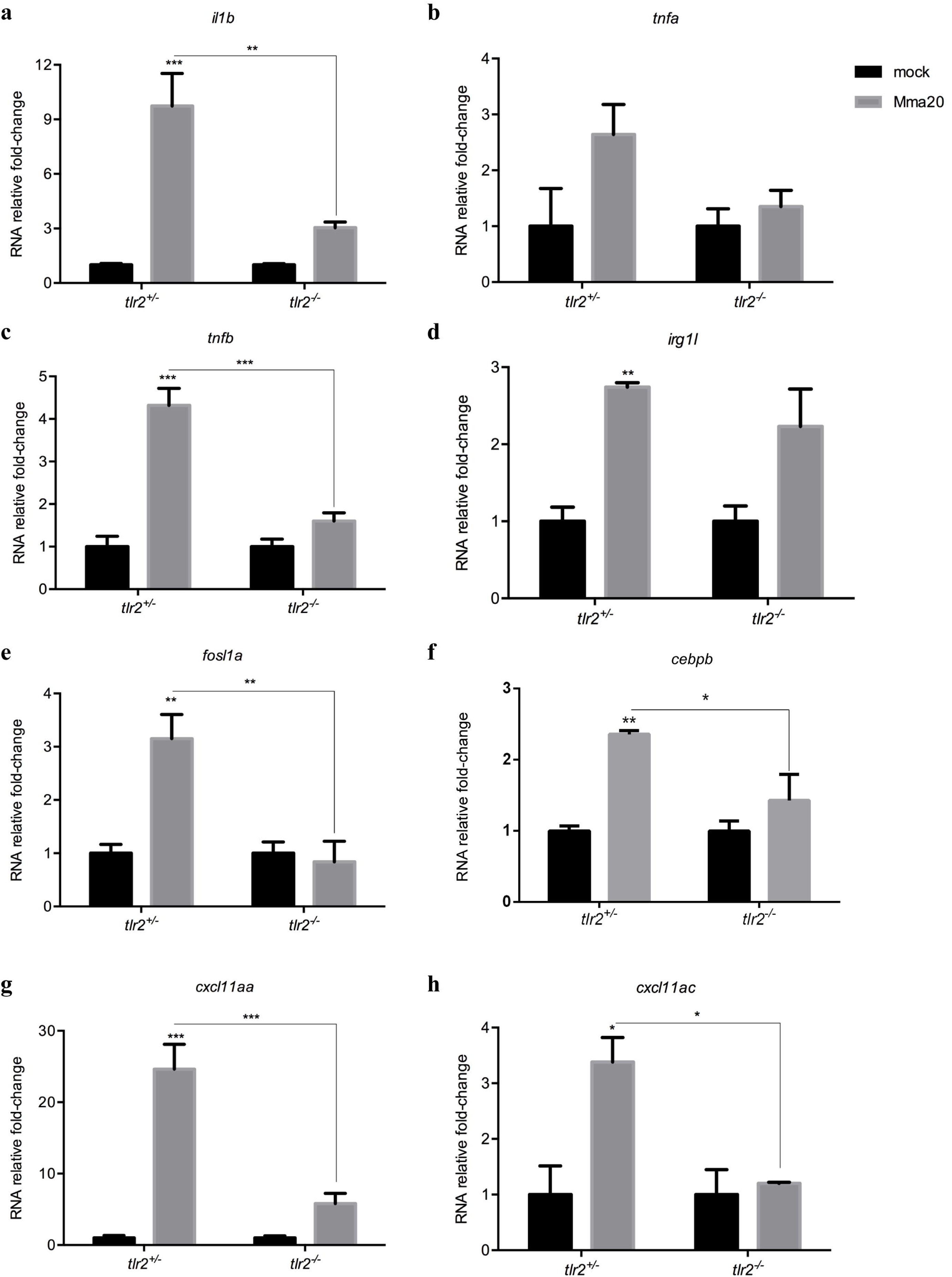
Immune genes expression in *tlr2*^+/−^ and *tlr2*^−/−^ fish lines infected with Mm. The expression levels of *il1b* (a), *tnfa* (b), *tnfb* (c), *irg1l* (d), *fosl1a* (e), *cebpb* (f), *cxcl11aa* (g) and *cxcl11ac* (h) were determined at 4dpi by qPCR. Data (mean ± SEM) are derived from at least three biological replicates (n = 10 embryos per group) and expressed relative to their corresponding mock injection (PBS) control, which is set at 1. Statistical significance of differences was determined by two-way ANOVA with Šidák Multiple Comparison test as a post-hoc test. \**P*<0.05, \*\**P*<0.01, \*\*\**P*<0.001

To further study the function of *tlr2* in defense against mycobacterial infection, we performed RNAseq of *tlr2^+/−^* and *tlr2^−/−^* larvae at 4 dpi with *M. marinum* strain (strain Mma20) and PBS as control. We summarized the number of differential expressed genes (DEGs) according to *P-*value (Fig. 5a, b) and in volcano plots (Fig. S4). The number of DEGs in *tlr2^+/−^* infected with Mm was higher than those in *tlr2^−/−^* at any given *P-*value or false discovery rate less than 0.05, or any given fold-change with a *P-*value less than 0.05. These data also show that most of the genes downregulated by the Mm infection in the control remained unchanged in *tlr2* mutants (Fig. 5a, b). To further analyze these RNAseq data, we chose the genes with a threshold of a *P*-value less than 0.05 in *tlr2^+/−^* with Mm infection (1102 up- and 827 down-regulated genes, Fig. 5a). Then, for these genes, we calculated the fold-change ratio of *tlr2^+/−^* versus *tlr2^−/−^*, and genes with ratios greater than 2 or less than 0.5 were selected for further analysis (Fig. 5c). As a result, 97 and 92 genes were scored as *tlr2* specific up- and down-regulated genes, respectively. Next, we conducted GO analysis (Fig. 5d, e) showing that genes grouped into the immune system category are the most prominently deregulated (36%) in the whole *tlr2* up-regulated 97-gene set (Fig. 5d). Within this category we found genes involved in lysosome, chemotaxis, transcription regulation, diverse immunoglobulin domain-containing proteins (*dicps*) and other immune processes (Fig. 5d and 6a-e). For other categories, many up-regulated genes fell in the categories oxidation-reduction process, DNA repair, transcription regulation and apoptotic process regulation (Fig. S5). In the *tlr2* down-regulated 92-gene set, the immune related genes also were the largest portion (15%; Fig. 5e, 6f). Many of these genes are poorly annotated and include genes encoding cysteine proteases, a *nitr* gene, a *mafb* transcription factor gene and *hsp70*. Categories encompassing non-immune related genes are listed in Fig. S6. These results show that, upon infection with Mm*, tlr2* mutants show a dampened response of immune genes.

**Figure 5.**
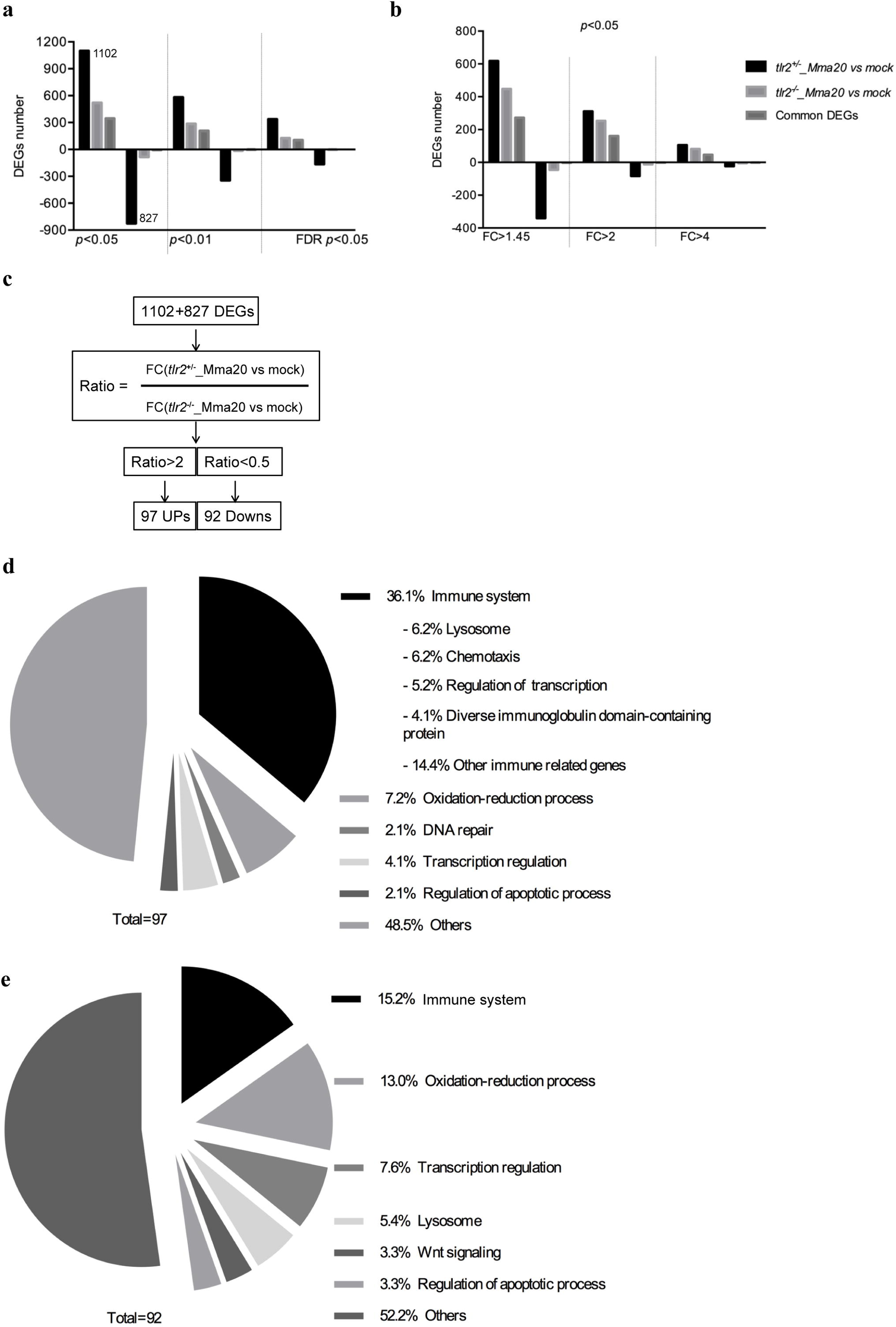
Overview of RNAseq results. (a, b) the number of DEGs of *tlr2*^+/−^ and *tlr2*^−/−^ strains infected with Mm compared to the control at different *p*-value and fold change. (c), the work flow of screening genes of which the regulation by infection is dependent on *tlr2*. (d), GO analysis of the 97 upregulated genes. (e), GO analysis of the 92 down-regulated genes. FC, fold-change.)

**Figure 6.**
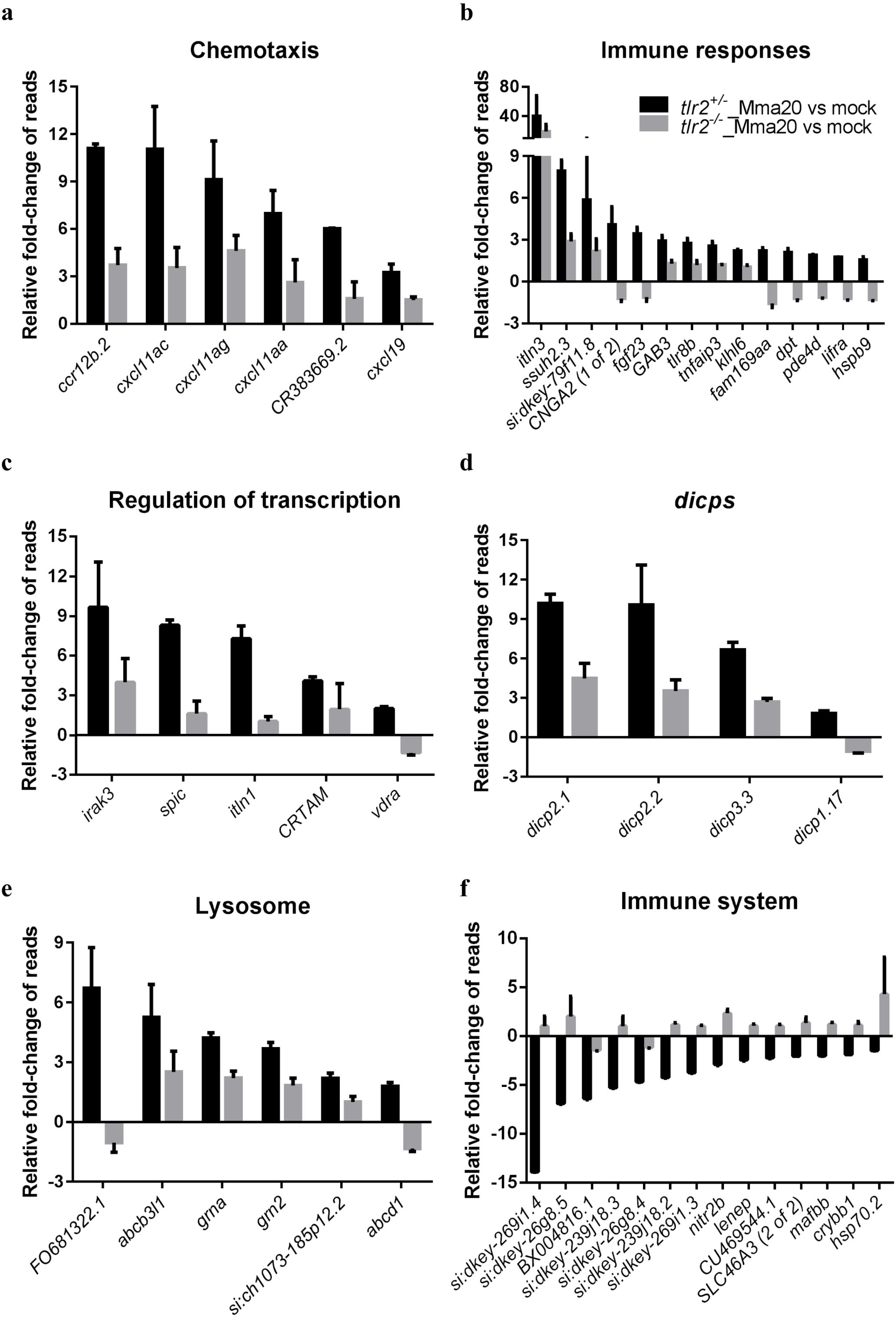
Overview of fold changes of representative genes selected from the gene categories resulting from GO-term analysis. (a-e): *tlr2*-dependent genes with up regulation corresponding to Fig. 5d. (f): *tlr2* specific genes with down regulation corresponding to Fig. 5e.

To show the relationship between DEGs, we constructed networks based on common expression targets in the 97 up- and 92 down-regulated genes [47]. The involved networks with the up-regulated genes contain *lipc*, *nr0b2*, *hmgcr*, *itln1*, *fgf23*, *vdr*, *irak3* and *tlr8* (Fig. S7). In *tlr2*^+/−^ controls we observed positively regulated *hmgcr*, *fgf23*, *vdr* and *tlr8*, and negatively regulated *nr0b2*, whereas *tlr2*^−/−^ showed an opposite regulation for most of these genes. However, *lipc* and *irak3*, were positively regulated in both *tlr2*^+/−^ and *tlr2*^−/−^. For the 92 downregulated genes, a network containing *nr0b2* and *mafb* was constructed (Fig. S8). *nr0b2* and *mafb* were positively regulated in *tlr2*^−/−^ zebrafish and down regulated in *tlr2^+/−^*.

### 4 Enrichment analysis of Tlr2 specific genes after Mm infection

To link our data to gene sets defined based on prior biological knowledge (including GO), we conducted Gene-Set Enrichment Analysis (GSEA) of the differently expressed genes. This method derives its power by focusing on gene sets, that is, groups of genes that share common biological function, chromosomal location, or regulation [48]. Because the number of predicted gene sets were too large for our analysis method (more than 1,000 with *P* < 0.05), we focused on the gene sets related to metabolic, immunological and inflammation pathways. GSEA predicted 61 pathways in *tlr2*^+/−^ and 67 pathways in *tlr2*^−/−^ zebrafish responsive to Mm infection (Supplementary Table 4A and 4B). Interestingly, most of these pathways are common in *tlr2*^+/−^ and *tlr2*^−/−^, however some of them are detected as being anti-correlated in regulation, including some pathways underlying natural killer cell functions and omega-6-fatty acid metabolism (Supplementary Table 4C). We also performed Sub-Network Enrichment Analysis (SNEA) [49] to identify possible key genes that are responsible for the difference in response of the *tlr2*^+/−^ and *tlr2*^−/−^ group to Mm infection (*P* < 0.05). SNEA predicted 565 and 503 pathways for *tlr2*^+/−^ and *tlr2*^−/−^ zebrafish that are linked to the response to infection, respectively (Supplementary Table 4D and 4E), and 264 and 202 of them are specific for the response in *tlr2*^+/−^ and *tlr*2^−/−^ fish, respectively (Supplementary Table 4F). Since the RNAseq was conducted with the total RNA from whole body of zebrafish, it is not possible to define which pathways are involved in macrophage functions related to *tlr2* expression. Therefore, we analyzed previously published DNA microarray data (GDS4781) of human macrophages transfected with *Mtb* from Gene Expression Omnibus [50]. SNEA analysis of human macrophages shows that 659 pathways are linked to Mtb infection (Supplementary Table 4G). By comparing the human SNEA result to *tlr2*^+/−^ specific pathways in zebrafish, 56 pathways were defined as *tlr2*^+/−^-specific (Supplementary Table 4H). Of these, the pathway of TLR8, which has the lowest *P-*value in both zebrafish and human enrichment (Supplementary Table 4H), and its network is depicted as example in Figure S9. Overall, these analyses show that *tlr2*^−/−^ mutants have a strongly altered immune response of which many pathways that are linked to human tuberculosis are differently responding.

## Discussion

TLR2 has been shown to play a role in host defense against Mtb in several rodent studies but its role in host innate immunity during infection is still not clear. Moreover, little is known about the systemic regulation of down-stream signaling of TLR receptors in animal models. As part of this study we generated a *tlr2* zebrafish mutant to study Tlr2 function in the innate immune system during mycobacterial infection. Our results show a clear effect of Tlr2 in defense against mycobacterial infection. The deep sequencing data provided the possibility to study the whole transcriptome profile in our mycobacterial infection model at the systems level. There are only a few RNA sequencing results of tuberculosis studies in rodents [51, 52], human serum [53], human pulmonary epithelial cells [54], and bovine systems.

To characterize the effect of *tlr2* mutation in the absence of infection, we compared the transcriptome of homozygous mutant larvae with that of heterozygote larvae, thereby excluding the effect of non-dominant background polymorphisms that might have resulted from ENU mutagenesis. The results show large differences between these genotypes, for instance in genes involved in metabolism (Fig. S3). In agreement, our previous results suggested a role in metabolism also for Myd88, the adaptor in TLR signaling [43]. The largest category of genes that was significantly affected in *tlr2* homozygous mutants was “neurological system process” (Supplementary table 2). Many recent studies show that a mutation in Tlr2 in mice resulted in effects of neuronal development and responses to injury [55, 56]. Some of these studies show a connection of Tlr2 deficiency to neuronal defects that could be related to Il10 function and autophagy [57, 58]. When focusing on the signaling pathways that could be involved, we observed that there was a significant effect in the GO category of transcription factors, namely the c-Maf factors that totals up to 546 representatives that were affected (Supplementary Table 3). The members of the Maf family of transcription factors, c-Maf and Mafb are specifically expressed in monocyte and macrophage lineages [59, 60], and in addition, c-Maf is also expressed in T helper cells [60]. c-Maf was also reported to directly regulate IL-10 expression induced by LPS in macrophages [61]. Double deficiency of Mafb/c-Maf promotes self-renewal of differentiated macrophages [62]. We did not detect differences in macrophage numbers between *tlr2* mutants and controls, but it remains to be further investigated if macrophage differentiation is altered as a result of Tlr2 deficiency.

Recently, we have shown that Tlr2 and its adaptor MyD88 are important for the response of the zebrafish host to the microbiome [43]. Therefore, it is very well possible that the transcriptional differences we find in the *tlr2* mutant versus the controls are due to an aberrant response to the microbiome. Similarly, we have found that mutation in *myd88* leads to a large difference in the zebrafish gene expression profile, even in the absence of pathogenic infection[43]. This difference appeared to be dependent on the presence of a microbiome. In future work we therefore aim to study the transcriptome response of the *tlr2* mutant under germfree conditions and in the presence of a microbiome under gnotobiotic conditions and investigate whether the dysregulation of neural development pathways in *tlr2* mutants might be linked to control of the gut-brain axis [63-67]. Such studies could also show whether the composition of the microbiome affects the progression of mycobacterial infection.

The analysis of differential gene expression during infection showed that there is a very pronounced effect of mutation of the *tlr2* gene. With respect to the number of genes affected, the strongest effect was observed in genes that are down regulated during infection, since this category was strongly diminished in the *tlr*2 mutant (Fig. 5a, b and Fig. S4). This observation suggests that Tlr2 has an important function in anti-inflammatory responses, in line with previous reports of studies in mice that showed a strongly decreased anti-inflammatory response in *Tlr2* knockouts [68, 69]. In a further analysis of the quantitative effects on the differences in expression levels of genes in both the up regulated and down regulated groups, we selected a number of genes that were most significantly affected in GO analysis (Fig. 5d, e). This GO analysis showed many groups to be affected with the immune response as the biggest group. We also performed GSEA and SNEA analyses (Fig. S7, S8, S9 and Supplementary Table 4) showing that the *tlr2* mutant has a very different immune response than heterozygote controls. Furthermore, we show that many signaling pathways that have been reported to be linked to tuberculosis infection in humans are differentially regulated in our data set. Most significantly, activation of the Tlr8 pathway was strongly affected (Fig. S9). This evidence suggests that the Tlr2 signaling is strongly connected with Tlr8 function. *TLR8* mutations (polymorphisms) increase susceptibility to mycobacteria in the human population [70, 71]. In addition, our analysis revealed differential expression of three other interesting categories of immune genes, discussed below.

The vitamin D receptor pathway genes that are normally up-regulated during infection in zebrafish larvae were down regulated in the *tlr2* mutant. Furthermore, pathway analysis (Fig. S7 and S9) also implicated the expression of the Tlr8 pathway connected to vitamin D signaling as being strongly affected in the *tlr2* mutant. Vitamin D has been shown to be an important regulatory factor during tuberculosis [72] and has been linked previously to TLR2 function in studies in cell cultures [73]. Therefore, aberrant vitamin D signaling could be a major contributing factor to the hyper-susceptibility phenotype of *tlr2* mutants in Mm infection.

Tlr2 has been shown to be essential for the up-regulation of a group of genes that encode the Diverse Immunoglobulin Domain-Containing Proteins (DICPS) (Fig. 5d). This group is as novel multigene family encoding diversified immune receptors. Haire *et al*. [74] reported that recombinant DICP Ig domains bind lipids and lipid extracts of different bacteria, including Mtb and Mm, a property shared by mammalian CD300 and TREM family members. In the down-regulated set also several DICP members appear to be dependent on Tlr2, such as *dicp1.17*, *dicp3.3*, that are linked to the GO term insulin-like growth factor binding. These correlations might relate to functions of Tlr2 in other processes such as the control of diabetes type II by gut microbiota. However, the DICP gene family lacks easily recognizable genetic homologs in mammals, making a translation to a function in mammalian tuberculosis and other diseases currently not yet possible [75].

Another highly relevant category of genes of which the induction or repression during infection is dependent on Tlr2 includes the chemokines. In a previous study of our laboratory, Torraca *et al*. demonstrated the function of the Cxcr3-Cxcl11 axis in macrophage recruitment to infection foci and showed that disruption of this axis by *cxcr3.2* mutation increases the resistance to mycobacterial infection [45]. Furthermore, by Fluorescence-activated cell sorting (FACS) of Mm-infected cells, we recently showed that *cxcl11a* is a robust marker of infected macrophages [46]. The infection induced expression of this chemokine is dependent on Myd88, the common adaptor of the majority of Tlrs, including *tlr2* [46]. In agreement, the *tlr2* mutant shows a significant lower expression of *cxcl11aa* and also of an related chemokine, *cxcl11ac*, during Mm infection (Fig. 4). Considering the large number of other chemokines that are controlled by Tlr2 during infection, it is clear that the integrative network of connections cannot yet be understood from these expression studies and need more detailed functional analyses, e.g. by combinations of different mutations or directed studies on responses to chemokines as shown by Torraca *et al* [45]. However, we can state that the phenotype of the *tlr2* mutant, at the infection level, and the level of transcriptional control such as the mentioned effects on regulation of MafB/c-Maf and chemokines shows a clear connection with macrophage chemotaxis. These evidences do not exclude that Tlr2 has many other functions during infection such as phagocytosis. For instance, Blander *et al*. [76] and Rahman *et al*. [77] showed that phagocytosis of bacteria and phagosome maturation are impaired in the absence of TLR signaling. Therefore, the large number of unannotated genes of which the expression during infection is dependent on Tlr2 is also worth studying in more detail in future studies.

## Conclusion

Our study shows that Tlr2, as a part of innate immunity, plays an important role in controlling mycobacterial infection as observed on the transcriptome and infection level. This function may be mediated by several mechanisms, including a general attenuation of the inflammatory response, reduced mycobacterial dissemination by dampening of CXCR3-CXCL11 signaling, and anti-mycobacterial effects like vitamin D signaling. Our results show that Tlr2 is a major Tlr family member upstream of Myd88 that activates the CXCR3-CXCL11 signaling axis. The *tlr2* mutant is therefore a valuable model for further studies using published infection models for other pathogens and the study of the interactions with gut microbiota in zebrafish larvae.

## Methods

### Zebrafish husbandry

All zebrafish were handled in compliance with the local animal welfare regulations and maintained according to standard protocols (zfin.org). Larvae were raised in egg water (60 g/ml Instant Ocean sea salts) at 28.5 °C. For the duration of bacterial injections, larvae were kept under anesthesia in egg water containing 0.02% buffered 3-aminobenzoic acid ethyl ester (Tricaine, Sigma-Aldrich, the Netherlands). The culture of zebrafish with mutations in immune genes was approved by the local animal welfare committee (DEC) of the University of Leiden (protocol 14198). All protocols adhered to the international guidelines specified by the EU Animal Protection Directive 2010/63/EU.

The *tlr2*^sa19423^ mutant line (ENU-mutagenized) was obtained from the Sanger Institute Zebrafish Mutation Resource (Hinxton, Cambridge, UK) and shipped by the Zebrafish Resource Center of the Karlsruhe Institute of Technology. The mutant allele was identified by sequencing. Heterozygous carriers of the mutation were outcrossed three times against wild type (*AB* strain), and were subsequently incrossed three times. Heterozygous fish of the resulting family were used to produce embryos. Homozygous mutants were outcrossed to the *Tg*(*mpeg1:mCherry-F);TgBAC(mpx:EGFP)* double transgenic line [78, 79], and the offspring with GFP and mCherry fluorescence were subsequently incrossed to produce the *Tg*(*mpeg1:mCherry-F);TgBAC(mpx:EGFP)* line.

### Bacterial strain preparation

The bacterial strain, *Mycobacterium marinum* m20 (Mma20) expressing mCherry fluorescent protein [80], was used in this study. For the infection to zebrafish larvae, the bacteria were prepared as previously described [81]. The infection inoculum was prepared in 2% polyvinylpyrrolidone40 solution (Calbiochem, the Netherlands), and 150 colony-forming units (CFU) of bacteria were injected into the blood stream at 28 hours post fertilization (hpf) as previously described [82].

### Ligands injection

Purified Pam3CSK4 (InvivoGen, France) and flagellin from *S. typhimurium* (Flagellin FliC VacciGrade™, Invitrogen, France) were diluted in 1 mg/ml and 100 μg/ml in sterile water, respectively. For injection, 1 nl of the ligand solutions were injected into the blood stream at 28hpf. Sterile water was injected as a control experiment. Injections were performed using a FemtoJet microinjector (Eppendorf, the Netherlands) equipped with a capillary glass needle.

### Bacterial burden imaging and quantification

Pools of 20 larvae were collected at 3- and 4-day post infection (dpi) and imaged by using the Leica MZ16FA Fluorescence Stereo Microscope (Leica Microsystems, Wetzlar Germany) equipped with the DFC420C color camera (Leica Microsystems). Bacterial loads were analyzed using dedicated pixel counting software as previously described [83].

### Confocal laser scanning microscopy imaging and image quantification

Larvae (2 dpf) were embedded in 1% low melting point agarose (Sigma Aldrich), and image acquisition was performed by using a Leica TCS SP8 confocal microscope (Leica Microsystems) with 10 times objectives (N.A. 0.4). Acquisition settings and area of imaging (in the caudal vein region) were kept the same across the groups for macrophages and neutrophils number counting (Fig. 1 j, k) and pixel counting (Fig. S1). Experiments were performed in two independent series. Double fluorescent lines *tlr2^+/+^ Tg*(*mpeg1:mCherry-F);TgBAC(mpx:EGFP)* and *tlr2^−/−^ Tg*(*mpeg1:mCherry-F);TgBAC(mpx:EGFP)* were used for macrophages and neutrophils number counting. Macrophage and neutrophil cell counting was either performed manually or by using the plugin Find Maxima in Fiji (http://imagej.nih.gov/ij/docs/menus/process.html#find-maxima) (Fig. 1 j, k) using projections of the z-series. 25 individual larvae for each group were used for counting and representative images are given in Fig. 1h, i. With manual counting the z-series data was examined in cases when it was unclear whether the fluorescent pixels presented one or multiple cells. The result of manual cell counting (Fig. S10) is comparable with the result of automated cell counting using Fiji. Pixel counting of the double transgenic lines (Fig. 1S a, b) was performed using dedicated pixel counting software as well as previously described [83].

### RNA isolation, cDNA synthesis and qPCR

Total RNAs were extracted using TRIzol Reagent (Life Technologies) and purified using RNeasy MinElute Cleanup Kit (Qiagen, the Netherlands). The concentration and quality of RNAs were evaluated by NanoDrop 2000 (Thermo Scientific, the Netherlands). cDNAs were synthesized from 1 μg total RNAs and qPCR were performed by using the iScript™ cDNA Synthesis Kit (BioRad, the Netherlands) and iQ™ SYBR Green Supermix (BioRad) and normalized against the expression of *ppial* as a housekeeping gene [84]. Results were analyzed using the ΔΔ t method [85]. Primer sequences are described in Supplementary Table 5.

### Deep sequencing and data analysis

Triplicates of 10 larvae of *tlr2^+/−^* and *tlr2^−/−^* with PBS (as control) or Mma20 injection, were homogenized in 300ul of TRIzol reagent, and total RNAs were purified as described above. RNAseq was performed using Illumina Hi-Seq 2500 as previously described [86]. The raw data is available in the NCBI GEO database under accession number GSE102766. The RNAseq data were mapped on the zebrafish genome (version GRCz10) and tag counts were performed by Bowtie 2 using GeneTiles software (http://www.genetiles.com) [87]. Then, we performed normalization and gene expression analysis using the R package and DESeq2 [88]. After statistical tests, we performed further bioinformatics analyses Gene-Set Enrichment Analysis [48], Sub-Network Enrichment Analysis [49] and Pathway Enrichment Analysis [89]. For creating gene networks based on common regulatory targets, we used Pathway Studio 9.0 (Elsevier, Amsterdam, the Netherlands) as previously described [47].

Comparison of edgeR and DEseq2: EdgeR and DEseq2 differ mainly in the aspects of normalization and estimation of the dispersion parameters. Normalization in edgeR is done via the trimmed mean of M values, while in DEseq2 this is done by comparing each library with a virtual library based on the relative log expressions. The dispersion parameters in edgeR are estimated by empirical Bayes and are therefore shrunken towards the overall mean of the estimates. Dispersion in DEseq2 is estimated by taking the maximum of the individual dispersions and the mean trend of the dispersions. As a consequence, edgeR tends to be more sensitive to outliers, while DEseq2 is less powerful [44].

### Statistical analyses

Graphpad Prism software (Version 7.00; GraphPad Software, San Diego, CA, USA) was used for statistical analysis. All experiment data are shown as mean ± SEM. In immune gene expression in *tlr2^+/−^* and *tlr2^−/−^* fish lines infected with Mm (Fig. 4), statistical significance of differences was determined by two-way ANOVA with Šidák Multiple Comparison test as a post-hoc test. The other experiments were analyzed by using unpaired, two-tailed t-tests for comparisons between two groups and one-way ANOVA with Tukey’s multiple comparison methods as a post-hoc test for comparisons between more than two groups. (ns, no significant difference; \**p*<0.05; \*\**p*<0.01; \*\*\**p*<0.001; \*\*\*\**p*<0.0001).

## List of abbreviations

CFU: colony-forming units
CLSM: confocal laser scanning microscopy
DEGs: differential expressed genes
Dpf: days post fertilization
Dpi: days post injection
FDR: false discovery rate
FACS: Fluorescence-activated cell sorting
GSEA: Gene-Set Enrichment Analysis
Hpf: hours post fertilization
Hpi: hours post injection
Mm: *Mycobacterium marinum*
Mtb: *Mycobacterium tuberculosis*
NMD: non-sense mediated mRNA decay
qPCR: quantitative polymerase chain reaction
RNAseq: Deep sequencing of cDNA derived from polyA RNA
SNEA: Sub-Network Enrichment Analysis
TB: tuberculosis

## Declarations

## Acknowledgments

We thank all members of the fish facility team for fish caretaking. We would like to thank Lanpeng Chen and Gerda E.M. Lamers for assistance with confocal laser scanning imaging. We also thank Vincenzo Torraca for helpful discussions and the qPCR primers of *cxcl11-*like genes. The zebrafish *tlr2* mutant was obtained from the Sanger Institute Zebrafish Mutation Resource (ZF-MODELS Integrated Project funded by the European Commission; contract number LSHG-CT-2003-503496), also sponsored by the Welcome Trust [grant number WT 077047/Z/05/Z].

## Funding

S. Y. and W.H were supported by a grant from the China Scholarship Council (CSC). This funding body had no role in the design of the study and collection, analysis, and interpretation of data and in writing the manuscript

## Availability of data and materials

The raw data of the RNAseq experiments is available in the NCBI GEO database under accession number GSE102766.

## Authors’ contributions

SY and WH carried out all the experimental operations. SY, YS, and HPS performed RNAseq data analysis. SY, WH and HPS wrote the first draft of the manuscript. SY, WH and HPS participated in the design of the study and coordination. AHM and RMJ helped to revise the final manuscript. HPS conceived the study. MM assisted in statistical analysis of RNAseq data. All authors read and approved the final manuscript.

## Ethics approval and consent to participate

No animals were used for experimentation. Larvae for experiments were obtained from zebrafish lines that were handled in compliance with the local animal welfare regulations and maintained according to standard protocols (zfin.org). The breeding of adult fish was approved by the local animal welfare committee (DEC) of the University of Leiden. All protocols adhered to the international guidelines specified by the EU Animal Protection Directive 2010/63/EU for which larvae under the age of 5 days post fertilization are not considered test animals.

## Consent for publication

Not applicable.

## Competing interests

The authors declare that they have no competing interests.

## Figure legends

**Supplementary figure 1** Pixel count analysis for double transgenic lines *Tg*(*mpeg1:mCherry-F);TgBAC(mpx:EGFP)* of 2 dpf *tlr2*^+/+^ and *tlr2*^−/−^ embryos. a: mCherry reporter, b, EGFP reporter.

**Supplementary figure 2** RNAseq read counts of *tlr2* transcripts in heterozygotes (*tlr2^+/^*^−^) control versus *tlr2* mutant (*tlr2^−/^*^−^) larvae. RNAseq data comparing reads mapped to *tlr2* transcript (ENSDART0000012256). Heterozygotes and mutant reads are mapped to the entire length of the mutant transcript indicating that the mutant transcript is not subjected to nonsense mediated decay. Mutant data (MEAS) and heterozygotes (CTRL) data have been submitted to the NCBI gene expression Omnibus database, accession number is GSE102766.

**Supplementary figure 3 Analysis of differential expression of genes functioning in glycolysis and gluconeogenesis between uninfected *tlr2^+/−^* and *tlr2^−/−^*.** The red boxes represent up regulated genes (FC >2); blue boxes represent down regulated genes (FC<-2); yellow boxes represent the genes that are differentially expressed with a *P* value lower than 0.05; green represent not-significantly differentially expressed genes.

**Supplementary Figure 4 Volcano plots.** Volcano plots showing the significance cutoff applied to *tlr2*^+/−^ infected with strain Mma20 versus control with PBS (a) and *tlr2*^−/−^ infected with strain Mma20 versus control with PBS (b). In these volcano plots, the transcripts were considered significant (red) or non-significant (blue) by the conditions of |fold change| ≥ □1,45 and adjusted *P* value threshold □≤□0,05. Fold changes for each transcript was plotted on the X-axis against −log10 transformed p-values on the Y-axis.

**Supplementary figure 5** *tlr2*-dependent up regulation of genes with various GO terms.

**Supplementary figure 6** *tlr2*-dependent down regulation of genes with various GO terms.

**Supplementary figure 7** Sub-network enrichment analysis. Networks of common targets of the 97 up regulated genes (Fig. 5d) in *tlr2*^+/−^ (a) and *tlr2*^−/−^ (b) with Mma20 infection. Red represents up regulation, blue represents down regulation and grey represents genes for which no expression was detected.

**Supplementary figure 8** Sub-network enrichment analysis. The networks of common targets of the 92 down regulated genes (Fig. 5e) in *tlr2*^+/−^ (a) and *tlr2*^−/−^ (b) with Mma20 infection. Red represents up-regulation, blue represents down-regulation and grey represents genes for which no expression was detected.

**Supplementary figure 9** Sub-network enrichment analysis between zebrafish and human. The Tlr8 pathway in zebrafish (a) with Mm infection and human macrophages (b) with Mtb infection. Red represents up-regulation, blue represents down-regulation.

**Supplementary figure 10** Manual counting analysis for *Tg*(*mpeg1:mCherry-F);TgBAC(mpx:EGFP)* of neutrophils (a) and macrophages (b) in 2 dpf *tlr2*^+/+^ and *tlr2*^−/−^ embryos.

**Supplementary Table 1** Reads of *mpeg1.1* and *mpx* genes of the *tlr2*^+/−^ control and the *tlr2*^−/−^ mutant in RNAseq analysis.

**Supplementary Table 2** GO analysis of genes that have a different basal expression level in the absence of infection in the Tlr2 mutant versus the heterozygote control. Shown is the GO analysis of the group of 878 genes that were different with a FDR *P* value of 10^−10^ indicated in Fig. 2.

**Supplementary Table 3** Two categories of genes including the c-Maf transcription factor were significantly affected in the GO category

**Supplementary Table 4 A-H** Gene lists of GSEA and SNEA analysis

**Supplementary Table 5** List of primers

