## Supplementary figures and images for "RNA-seq analysis of a zebrafish *tlr2* mutant shows a broad function of this Toll-like receptor in transcriptional and metabolic control and defense to *Mycobacterium marinum* infection"

### Supplementary Fig 1

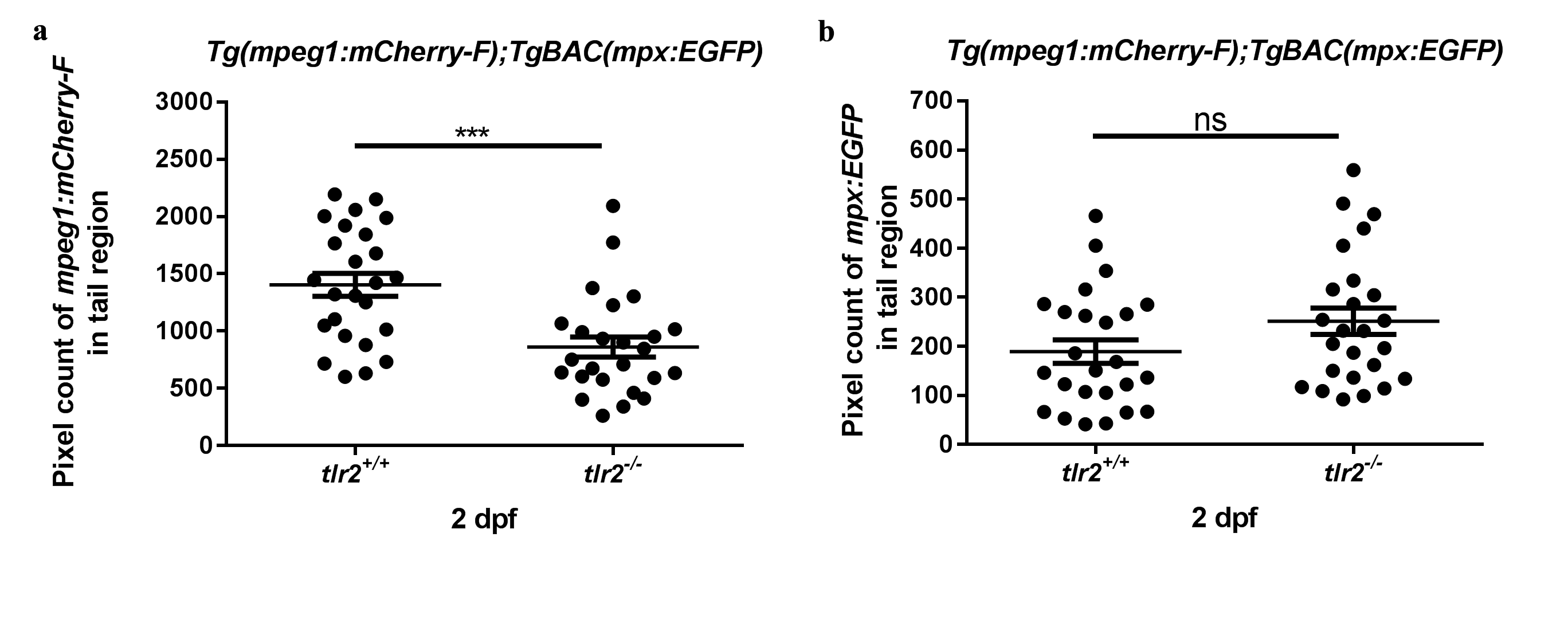

### Supplementary Fig 2

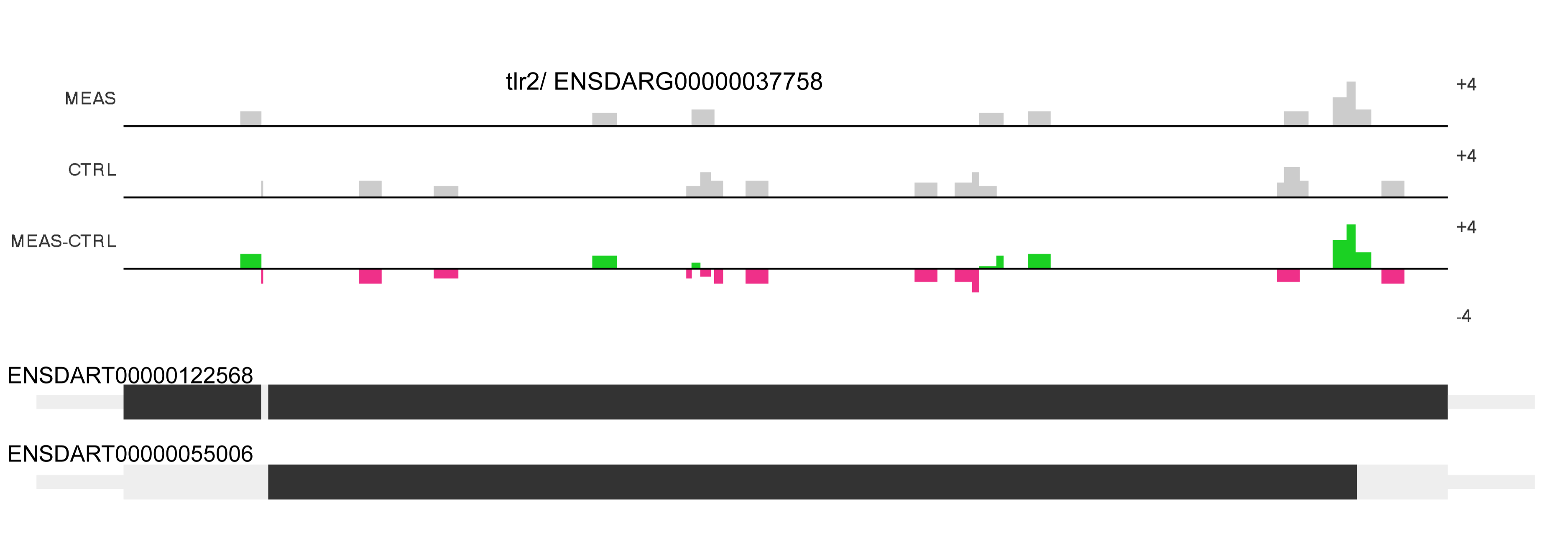

### Supplementary Fig 3

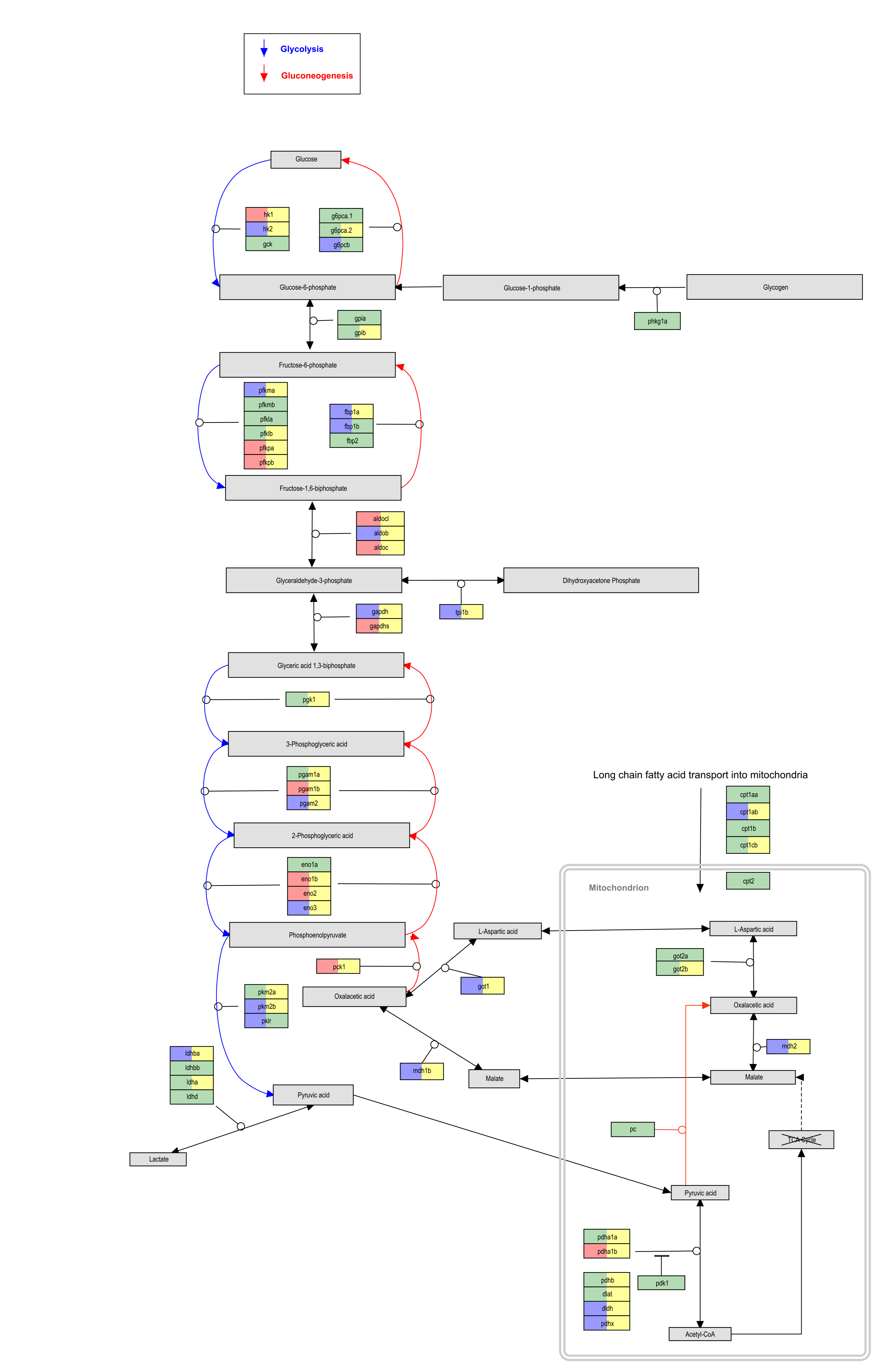

### Supplementary Fig 4

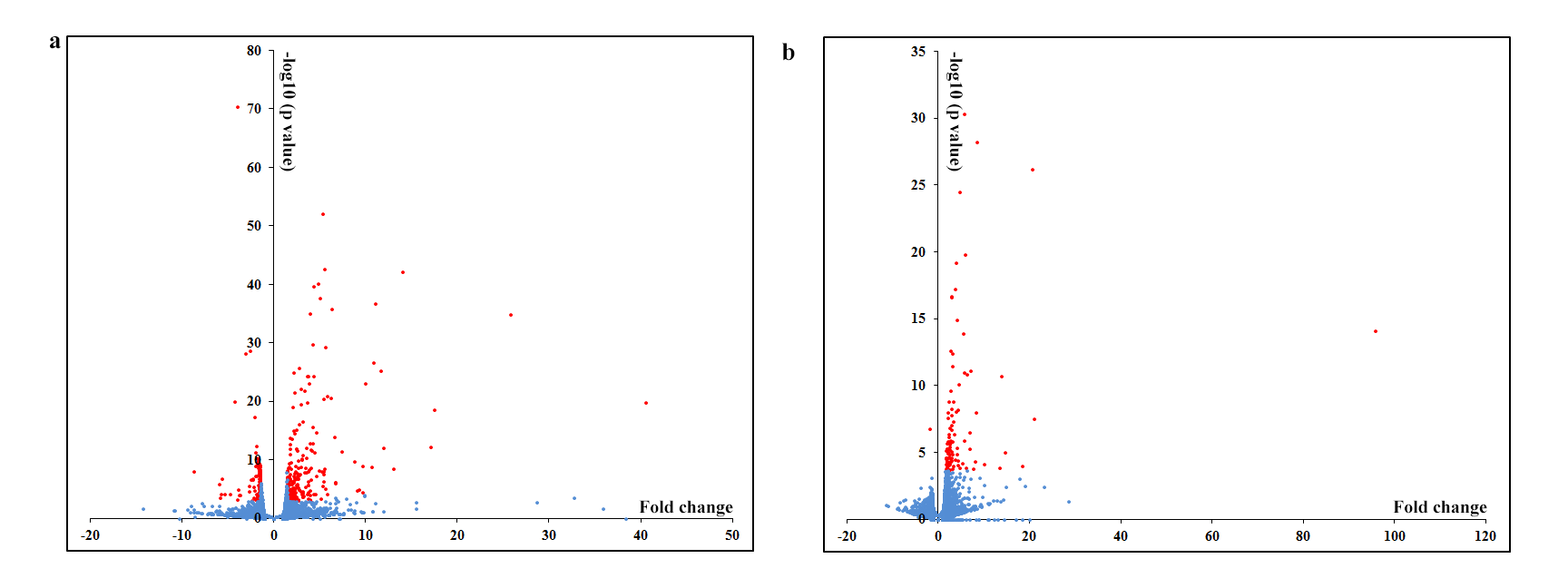

### Supplementary Fig 5

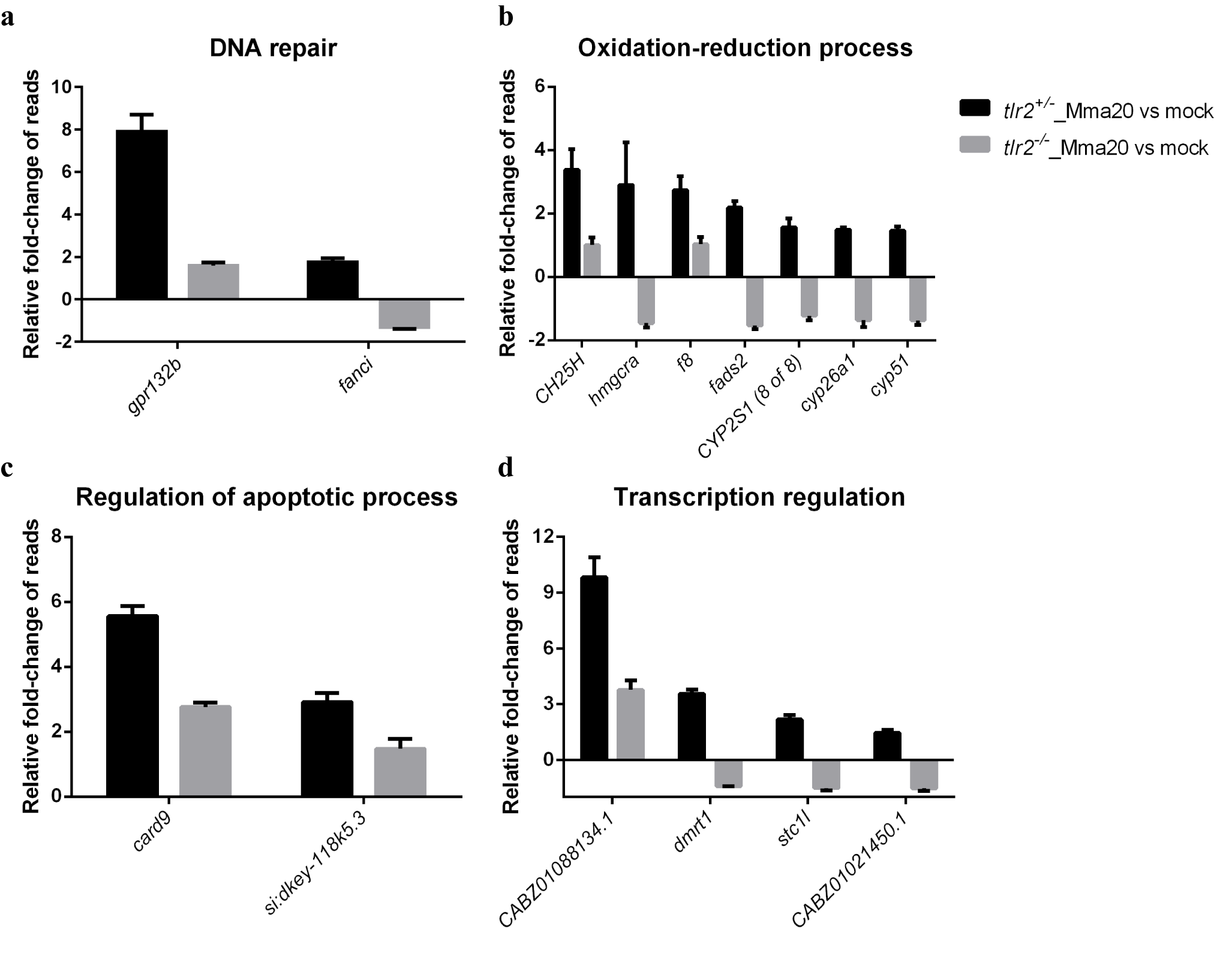

### Supplementary Fig 6

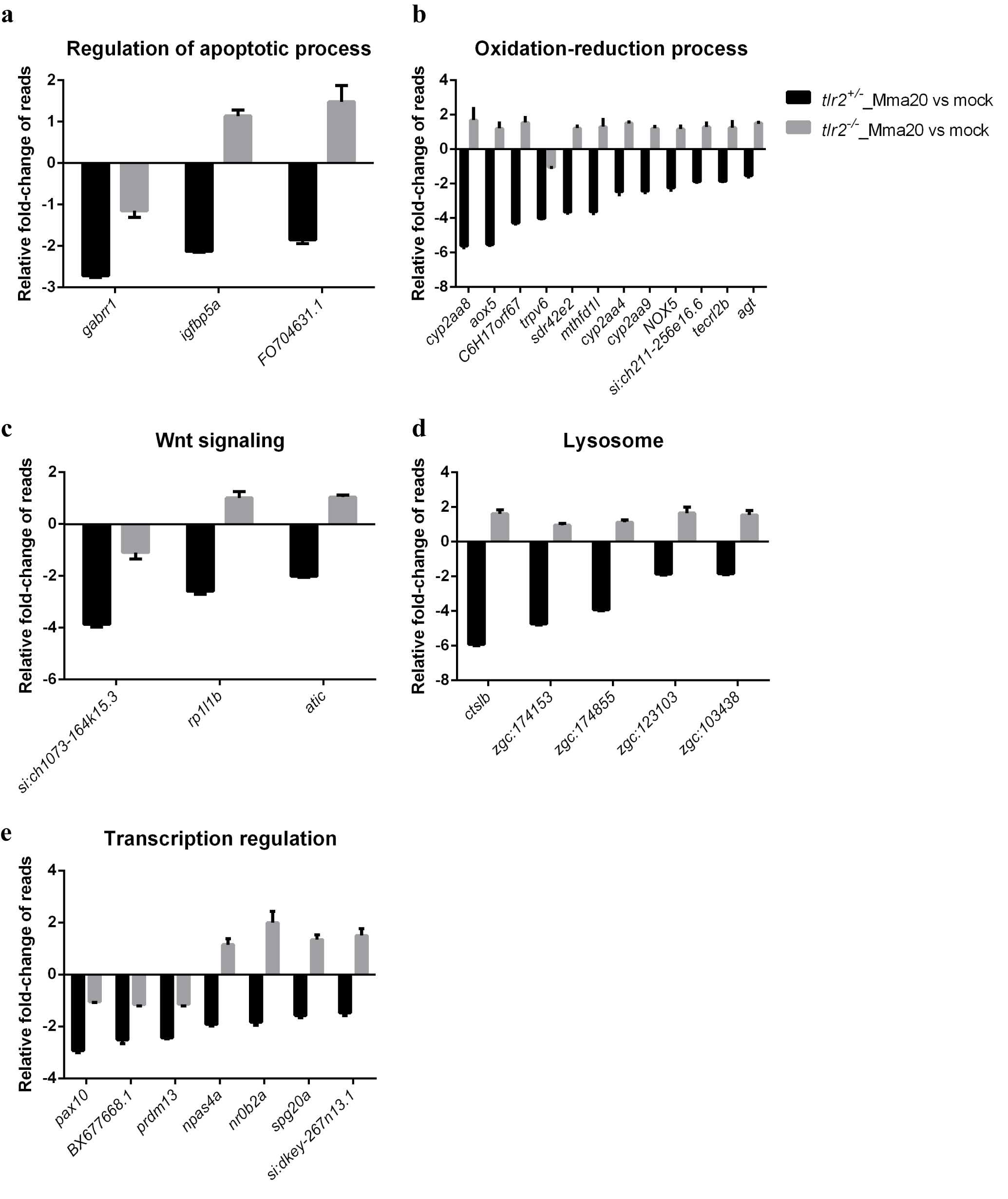

### Supplementary Fig 7a

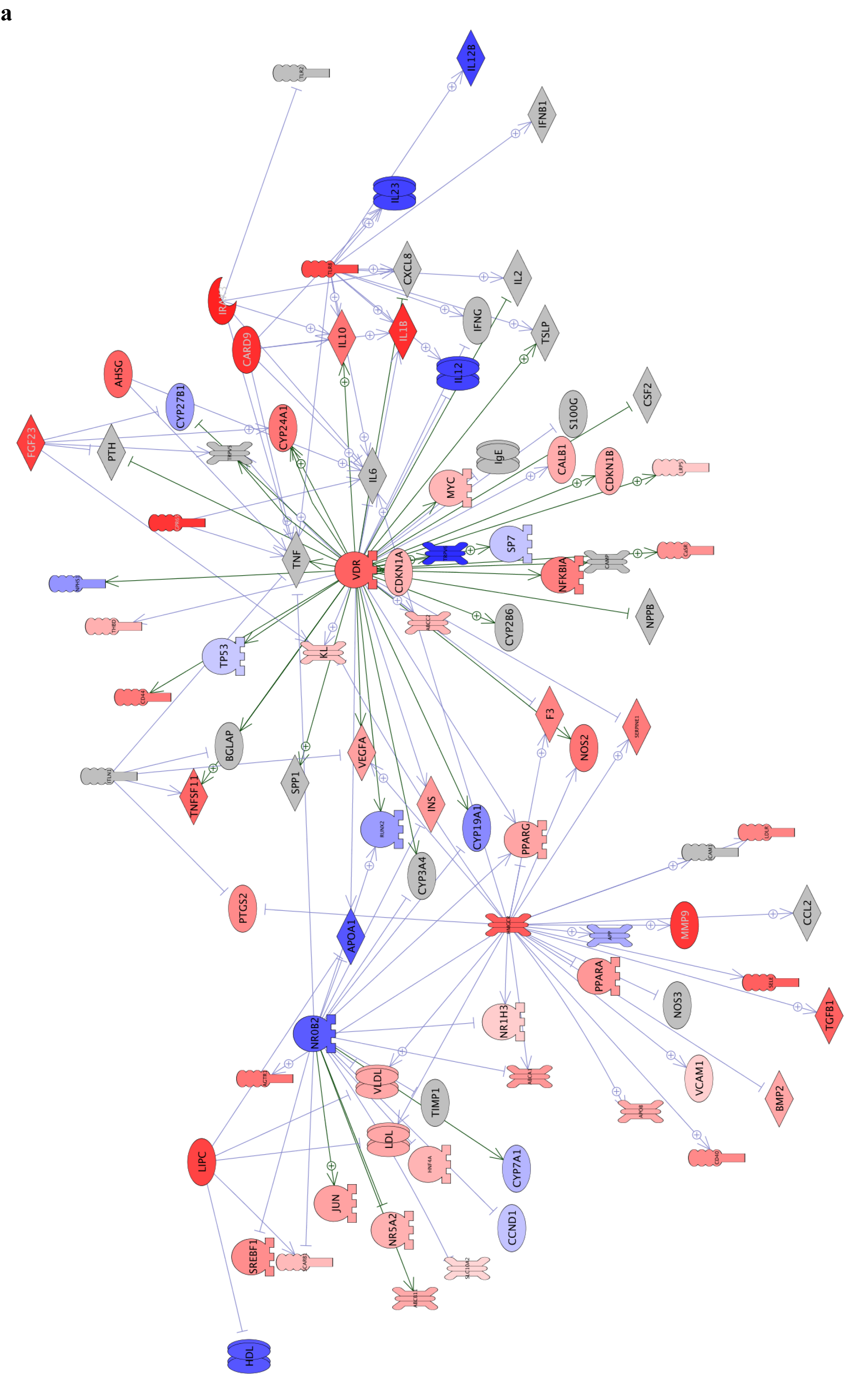

### Supplementary Fig 7b

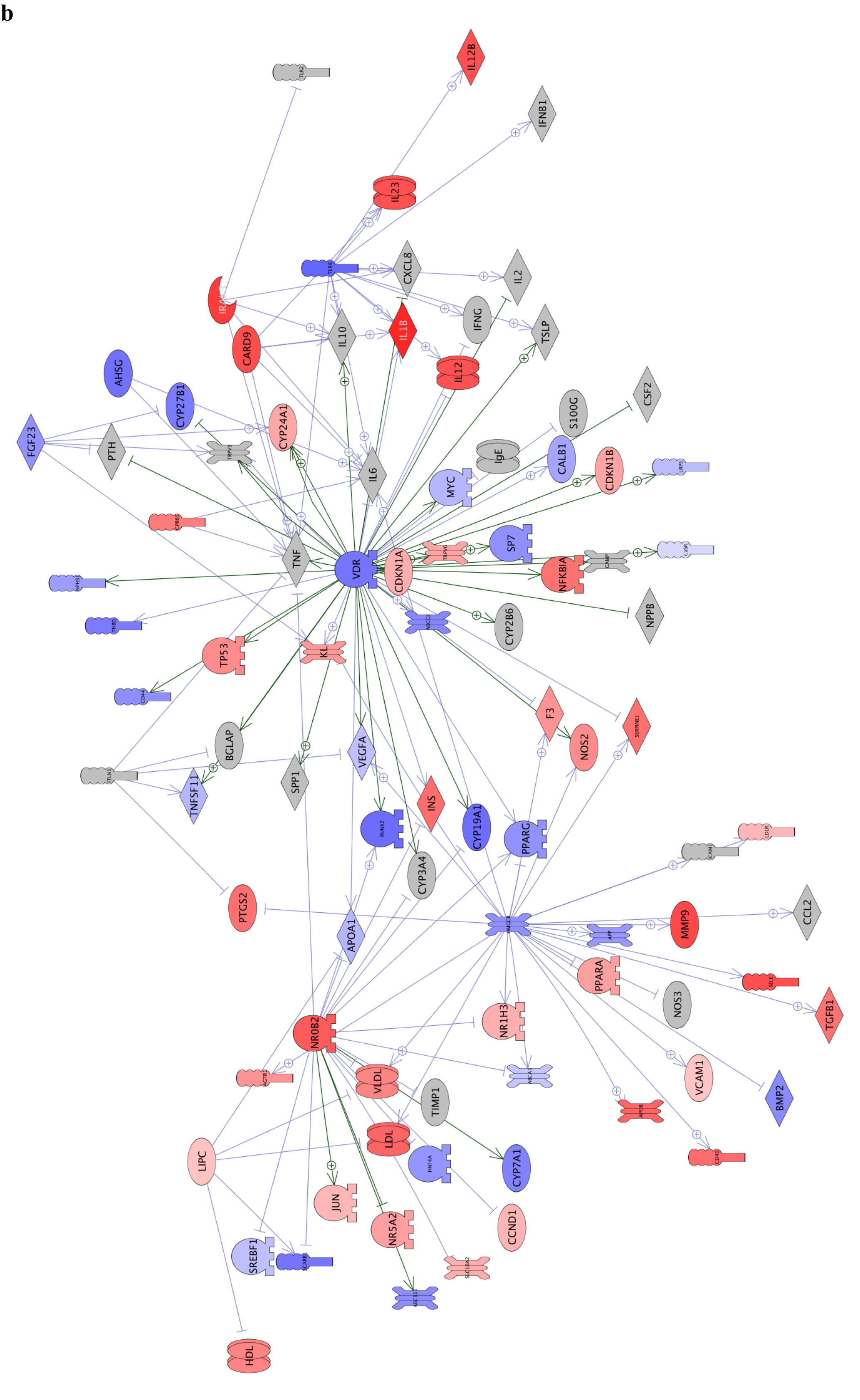

### Supplementary Fig 8

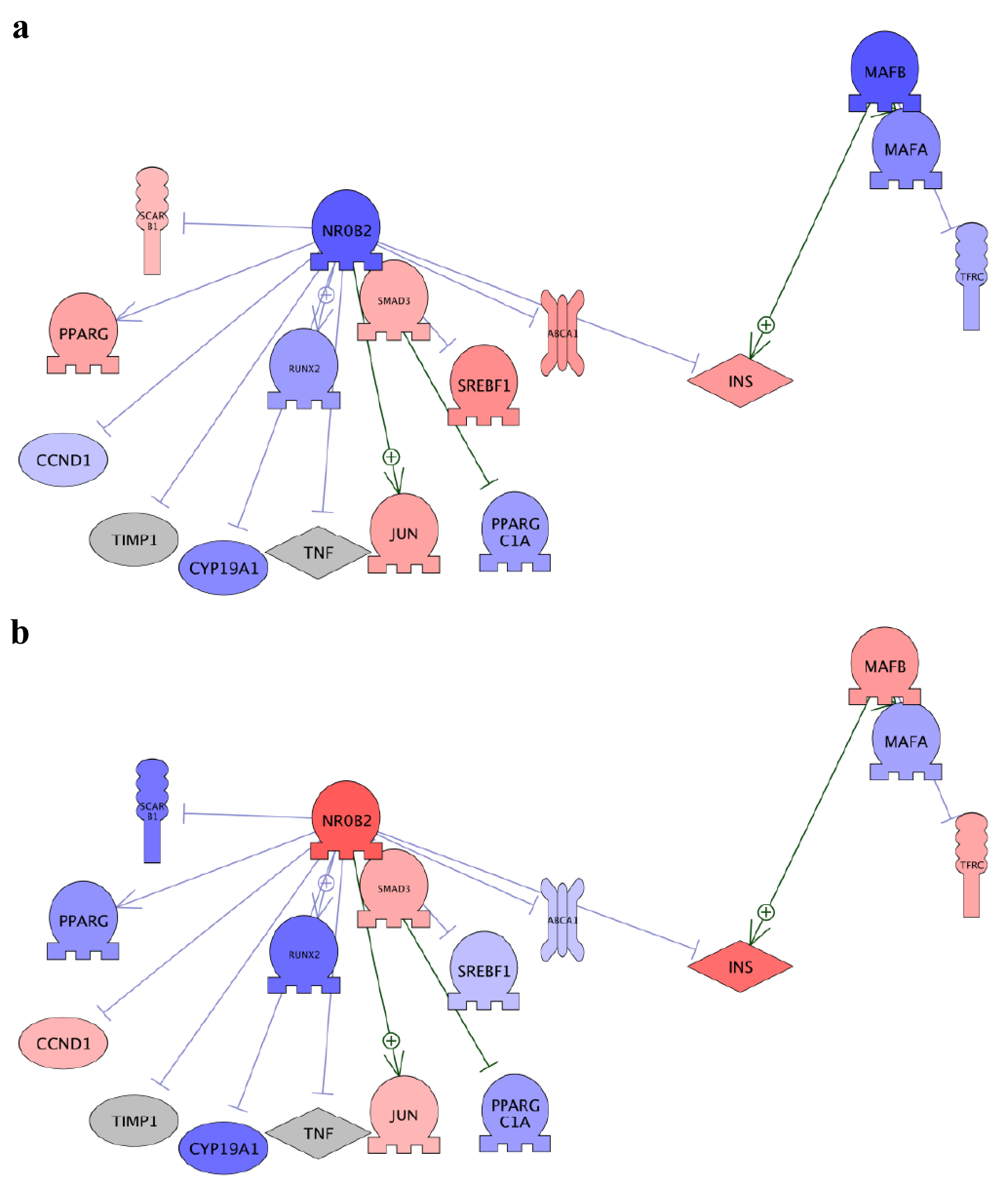

### Supplementary Fig 9

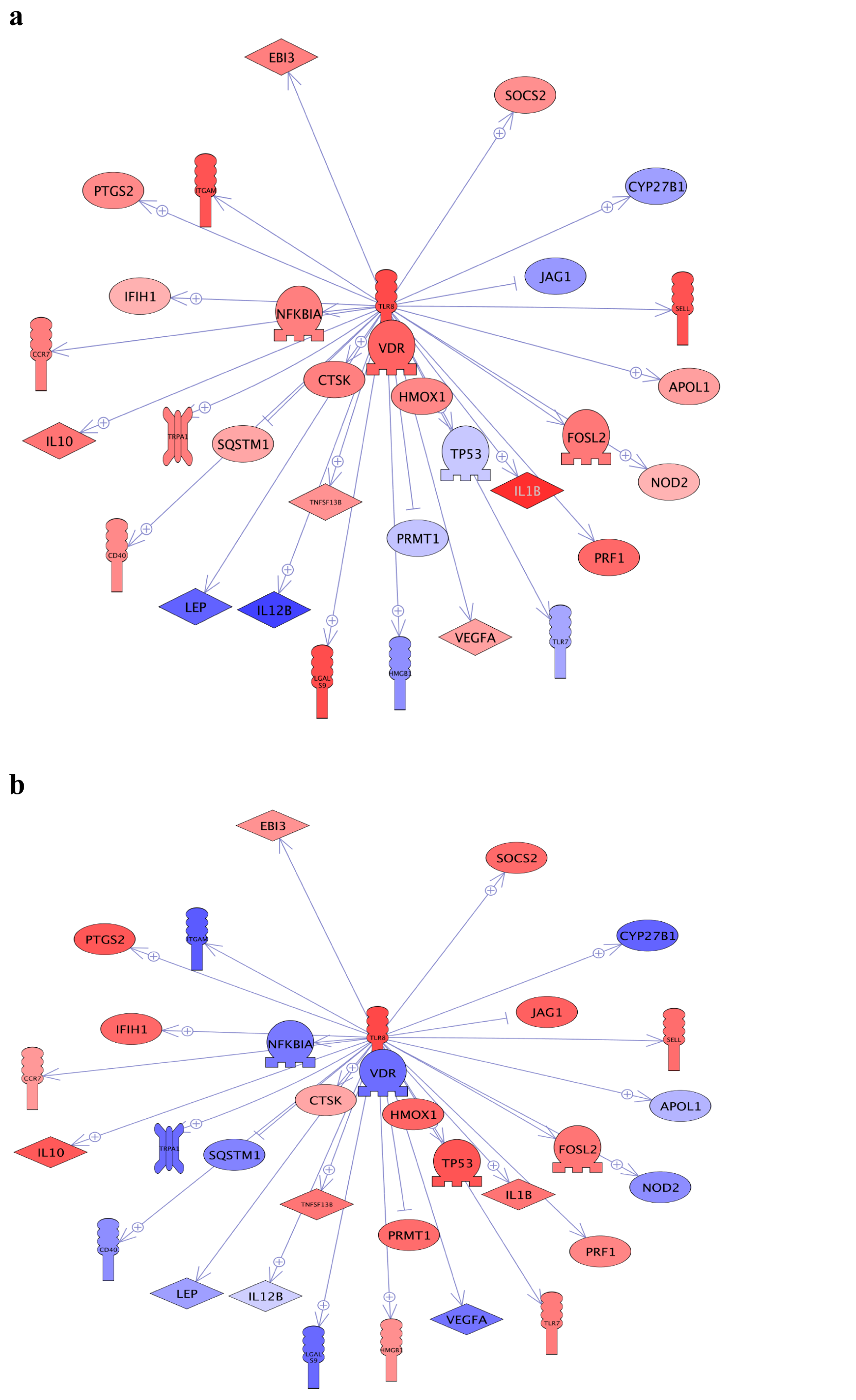

### Supplementary Fig 10

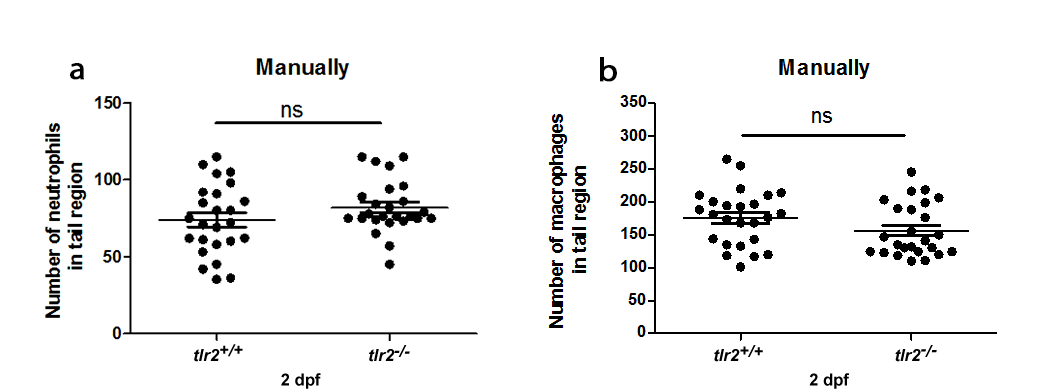
